# Hemoglobin in the diet modulates post-blood feeding behavioral rhythms and gene expression in *Aedes aegypti*

**DOI:** 10.1101/2025.08.29.673051

**Authors:** Diane F. Eilerts, Karthikeyan Chandrasegaran, Oluwaseun M. Ajayi, Ajay Sharma, Olivia Evans, Morgen VanderGiessen, Joshua B. Benoit, Clément Vinauger

## Abstract

Female *Aedes aegypti* mosquitoes rely on blood meals to acquire nutrients essential for egg development. However, blood ingestion also introduces physiological stressors—including thermal, osmotic, and oxidative stress—particularly during the digestion of heme-containing proteins. In *Drosophila melanogaster*, oxidative stress response genes are regulated by the circadian clock, and core circadian transcription factors are redox-sensitive. Additionally, sufficient iron ingestion is necessary to maintain normal circadian behaviors, as iron metabolism genes influence circadian behaviors, and flies lacking iron storage and transport are arrhythmic. However, whether similar interactions between iron metabolism and circadian rhythms exist in mosquitoes remains unclear. Here, we leverage the known alteration of mosquitoes’ activity rhythms following a bloodmeal to investigate whether dietary hemoglobin contributes to post-feeding activity suppression and gene regulation in the female *Ae. aegypti*. Using vertebrate blood and artificial blood mimic diets of equal nutritional values but with and without hemoglobin, we compared mosquito egg production, locomotor activity, sleep profiles, and transcript abundance of genes involved in circadian regulation and host-seeking behavior. Hemoglobin intake significantly reduced post-feeding activity, increased sleep amounts, and suppressed transcription of the core circadian gene *period* in mosquito heads. Short periods of sleep deprivation during the post-blood feeding period of inactivity did not alter egg production, timing of deposition, or viability. Our findings reveal that hemoglobin-derived heme influences behavioral and molecular responses in *Ae. aegypti* after a blood meal, pointing to a complex regulatory network linking heme acquisition, oxidative stress, daily rhythms, and behavior.

## 1. Introduction

Adult mosquitoes of both sexes obtain energy from plant-derived carbohydrates such as nectar, fruit juices, and honeydew (Barredo and DeGennaro, 2020; Foster, 1995; Upshur et al., 2023). These sugars support essential behaviors, including flight, reproduction, and host-seeking (Peach and Gries, 2020). However, females require a blood meal to acquire the proteins and nutrients necessary for oogenesis (Greenberg, 1951). Blood-feeding introduces the risk of pathogen acquisition, ultimately enabling disease transmission, while imposing thermal, osmotic, and oxidative stress resulting from the digestion of heme-containing proteins (Benoit et al., 2010; Benoit and Denlinger, 2017; Lahondère and Lazzari, 2012; Saeaue et al., 2011).

A major component of the bloodmeal is heme, which acts as the source of iron for the eggs (Zhou et al., 2007). As a pro-oxidant component of hemoglobin, heme can also generate reactive oxygen species (ROS) through the Fenton reaction, leading to cellular damage, lipid membrane disruption, and even cell death (Chaitanya et al., 2016). Blood-feeding mosquitoes have evolved multiple strategies to manage heme toxicity, including the formation of a peritrophic matrix that shields midgut tissues, and the induction of antioxidant and detoxification enzymes such as catalase, superoxide dismutases, and glutathione S-transferases (Oliver and Brooke, 2016; Saeaue et al., 2011; Whiten et al., 2018). Additionally, *Aedes aegypti* employs a unique heme degradation pathway in which heme oxygenase breaks down heme into biliverdin, carbon monoxide, and free iron, followed by glutamine conjugation of biliverdin (Pereira et al., 2007; Wilks and Heinzl, 2014). Thus, heme acquired from the bloodmeal is likely to have critical roles in both nutrition and in the generation of oxidative stress during blood digestion. Despite this complexity, the behavioral consequences of dietary heme remain poorly understood.

Nutritional levels and oxidative stress are likely to strongly influence behavioral and circadian processes. In *Drosophila melanogaster*, oxidative stress sensitivity disrupts circadian rhythmicity at both behavioral and molecular levels, with redox-sensitive transcription factors playing key roles in clock regulation (Hardeland et al., 2003; Krishnan et al., 2008; Mandilaras and Missirlis, 2012; Zheng et al., 2007). Food ingestion and nutritional status have a direct impact on daily and circadian rhythms (Hwangbo et al., 2023; Villanueva et al., 2019). This relationship between oxidative stress, nutrition, and circadian function in *Drosophila* raises the question of whether similar or more pronounced mechanisms are present in mosquitoes, where blood meals are nutritionally essential, particularly iron-rich, and have been documented to be rhythm-disrupting (Holmes et al. 2025; Dong et al. 2025).

In *Ae. aegypti*, a major arbovirus vector, locomotor activity exhibits diurnal peaks at dawn and dusk (de Lima-Camara, 2010; Eilerts et al., 2018; Kawada et al., 2005; Taylor and Jones, 1969). Almost symmetrically, *Ae. aegypti* mosquitoes enter phases of sleep-like states during the night and a few hours before midday (Ajayi et al., 2024, 2022, 2020). These rhythms are modulated by physiological states, including insemination, infection, and blood-feeding status (Evans et al., 2009; Gentile et al., 2013; Lima-Camara et al., 2014, 2011; Luz et al., 2011). Following a blood meal, mosquitoes exhibit suppressed host-seeking and locomotor activity, likely driven by abdominal distention and hormonal signals associated with blood digestion and oocyte maturation (Dong et al., 2025; Holmes et al., 2025; Klowden and Lea, 1979; Takken et al., 2001). Post-bloodmeal behavioral changes can be influenced by multiple factors, including incomplete blood ingestion (Duvall, 2019; Scott et al., 1993) and challenges in locating optimal resting sites (Holmes et al., 2025).

Although it is well-established that blood-feeding suppresses mosquito activity and host-seeking behavior, the role of heme or hemoglobin digestion in driving these behavioral changes remains unclear. It is also unknown whether heme-induced oxidative stress feeds back on central or peripheral circadian clocks to modulate behavioral rhythms in mosquitoes, as has been demonstrated in *Drosophila* (Mandilaras and Missirlis, 2012; Rudisill et al., 2019). To address this knowledge gap, we examined how dietary hemoglobin influences reproductive output, locomotor activity, sleep, and gene expression in *Ae. aegypti* females. Using both artificial membrane feeders and live-host feeding, we compared animal blood meals with hemoglobin-manipulated artificial diets to isolate the role of dietary heme. We quantified locomotor activity and transcript levels of genes associated with circadian regulation (*period*), host-seeking (foraging, neuropeptide Y-like receptor I, and short neuropeptide F), and oxidative stress management (catalase and heme oxygenase). Finally, sleep-deprivation assays evaluated whether post-bloodmeal sleep is necessary to maximize egg production.

## 2 Materials and Methods

### 2.1 Mosquito rearing

*Aedes aegypti* Liverpool (LVP-IB12, MR4, ATCC®, Manassas, VA, USA) larvae were raised in 26 × 35 × 4 cm covered trays filled with ~1 cm of deionized water, at a density of approximately 200 larvae per tray. They were maintained under light:dark (LD) cycles of 12 h:12 h at 26 °C and 70 ± 10% humidity. Larvae were maintained on daily feedings of Hikari Tropic First Bites (Petco, San Diego, CA, USA). Pupae were isolated on the day of pupation and placed into mosquito breeding containers (BioQuip, Rancho Dominguez, CA, USA). Females were maintained in the presence of males until being isolated for experiments to ensure they were mated and thus more motivated to host-seek and blood-feed.

For studies on sleep deprivation and live host feeding, mosquito larvae (Gainesville strain) were reared in plastic pans (30.5 × 7.5 × 5 cm) at a consistent density of ~200 larvae per pan. The rearing medium consisted of deionized (DI) water, finely ground fish food (Tetramin Goldfish Flakes), and yeast extract (Difco, BD 210929). Colonies were maintained in a climate-controlled facility (27 ± 2 °C; 65–70 % RH; 16:8 h light–dark cycle). Adults were housed in mesh cages (30.5 × 30.5 × 30.5 cm) with ad libitum access to water and 10 % sucrose solution. Females (12–14 days old) were blood-fed on a human host (University of Cincinnati IRB, 2021-0971) to sustain the colony and for studies post-blood feeding.

### 2.2 Artificial test diets

We used an artificial blood meal replacement diet (Gonzales et al., 2018) to explore the role of heme-containing proteins in the decreased activity after blood-feeding. For all experiments, mosquitoes were fed during their normal host-seeking peak, *i.e.*, the last two hours of their photophase (ZT 10–12), on one of four test diets: a 10% sucrose solution, heparinized bovine blood, an artificial protein diet containing hemoglobin (modified from (Gonzales et al., 2018)), and an artificial protein diet without hemoglobin (see **Table 1**). Only mosquitoes that had a visibly distended abdomen after feeding were used for oviposition, actometer, and transcript quantification experiments. The two artificial diets were prepared by dissolving components in an alkaline phosphate buffer according to (Gonzales et al., 2018) (see **Table 1** for final concentrations). Artificial diets will be referred to as “*Hb*” or “*noHb*” diet to differentiate the two artificial protein diets by the presence or absence of hemoglobin, respectively.

**Table 1.**
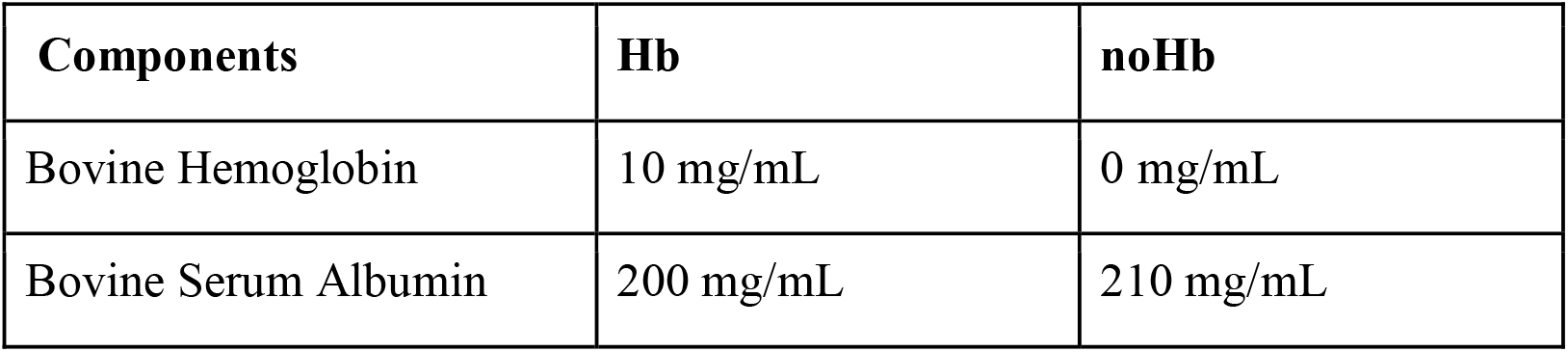
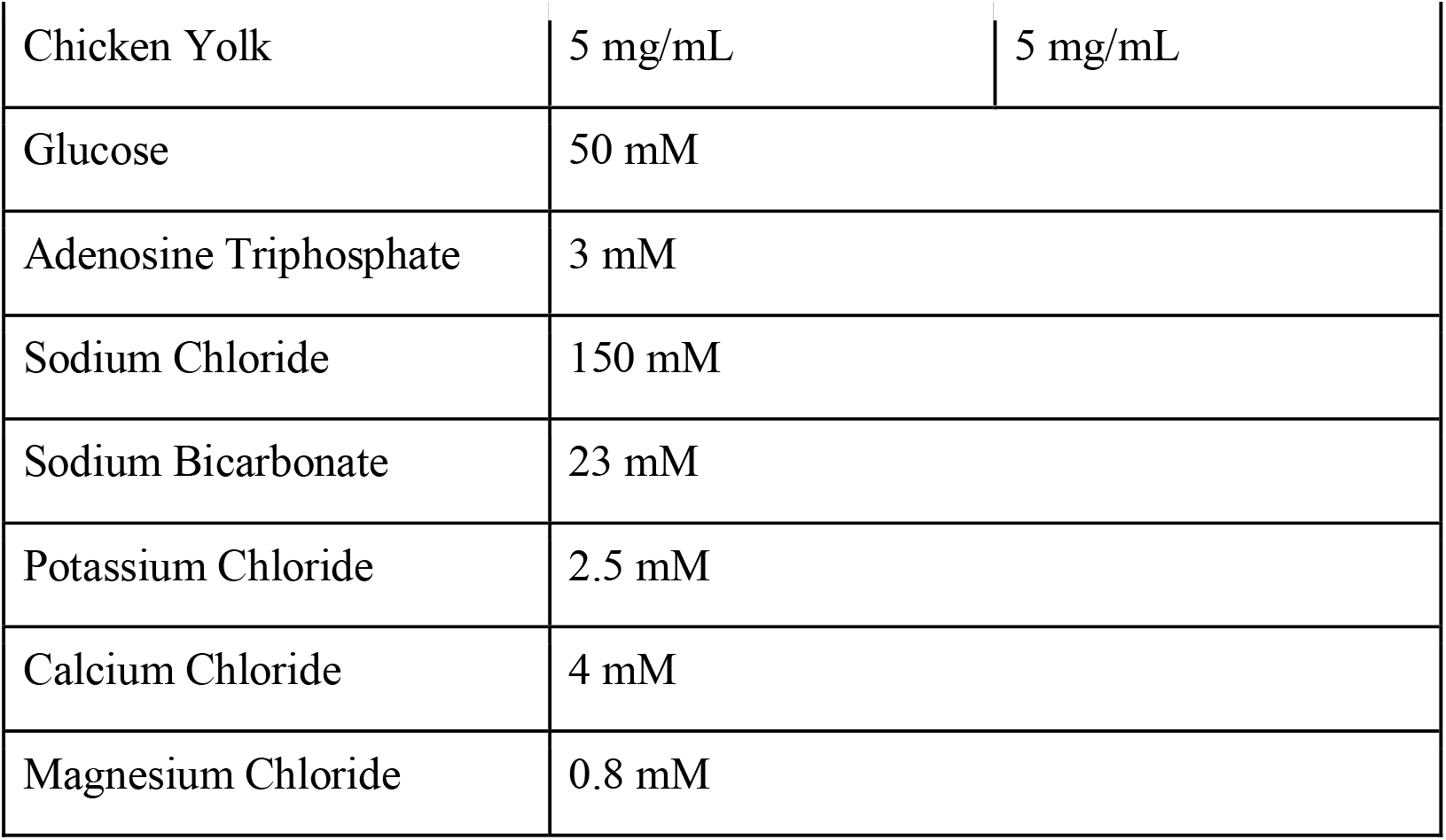
Components and final concentrations of the Hb and noHb diets. Modified from (Gonzales et al., 2018).

| Components | Hb | noHb |
| --- | --- | --- |
| Bovine Hemoglobin | 10 mg/mL | 0 mg/mL |
| Bovine Serum Albumin | 200 mg/mL | 210 mg/mL |
| Chicken Yolk | 5 mg/mL | 5 mg/mL |
| Glucose | 50 mM |  |
| Adenosine Triphosphate | 3 mM |  |
| Sodium Chloride | 150 mM |  |
| Sodium Bicarbonate | 23 mM |  |
| Potassium Chloride | 2.5 mM |  |
| Calcium Chloride | 4 mM |  |
| Magnesium Chloride | 0.8 mM |  |

The Hb and noHb diets contain the same total protein mass, with removed hemoglobin from the noHb diet replaced by excess bovine serum albumin (BSA), as shown in **Table 1**. Amino acid composition of all *Bos taurus* (bovine) hemoglobin subunits was compared to that of bovine BSA to confirm that all amino acids present in hemoglobin would still be available in BSA. Indeed, we found that amino acid frequencies were similar, and actually slightly higher in BSA for critical amino acids such as isoleucine and proline. Isoleucine is essential for oogenesis and reproduction, while proline supports post-blood meal physiology, ammonia detoxification, and energy production (Briegel, 1985; Rivera-Pérez et al., 2017; Scaraffia et al., 2008, 2005; Scaraffia and Wells, 2003).

### 2.3 Gut dissection

Upon feeding on bovine blood, the Hb diet, and the noHb diet, fully engorged mosquitoes were isolated, soaked in 1X Dulbecco’s Phosphate Saline Buffer (DPBS without Ca and Mg), and stored at 4°C for 45 minutes. Under a dissection microscope, individual mosquitoes were placed on a glass slide with their lateral side up and illuminated from below. The wings and legs were removed using forceps, following which 100 µL of Hank’s Balanced Salt Solution buffer (HBSS; Sigma H9394) was added to the sample. The HBSS buffer helps maintain osmotic balance and prevents the sample from drying out. The mosquito was dissected with a dissection needle, and a syringe needle (25G × 1^1/2^) was used to stabilize the sample. Dissection began with an incision in the lateral thorax above the hind leg. The incised cuticular tissue was held and further pulled apart downward until the last abdominal segment, exposing the ovaries, trachea, gut, malpighian tubes, and rectum. The cuticle on the medial side was ripped upward until the mesonotum, exposing the crop and salivary glands. The crop and the gut were retained for imaging, while the remaining tissue and scales were carefully removed. Mixing the Hb and noHb test diets with food coloring dyes aided in visualizing the consumed meal in the gut (Spice Supreme Green & Blue). The dissected gut and the crop were positioned as desired using a fine paintbrush and were imaged using a Microscope Digital Camera (AmScope MU1003).

### 2.3 Oviposition and egg viability assays

Mosquitoes aged 7–9 days were starved for 24 h and then provided either bovine blood, the Hb diet, or the noHb diet, warmed to 37 □via a glass artificial membrane feeder (Lillie Glassblowers, Smyrna, GA). Females that were visibly engorged after feeding were anesthetized on ice until inactive and then individualized into clear polystyrene *Drosophila* vials (25 × 95 mm Fly tubes, Genesee Scientific, San Diego, CA). Vials were prepared with a small water-laden cotton piece (about 2.5 cm) at the bottom, covered with a round piece of germination paper cut to the vial diameter, and enclosed with mesh. Mosquitoes were provided with a 10% sucrose-soaked cotton ball placed over the mesh, which was replaced daily. Individualized mosquitoes were maintained under light:dark (LD) cycles of 12 h:12 h at 26 ± 1 °C and 70 ± 10% humidity. Over the course of 9 days, mortality was recorded daily, and oviposition was monitored. Egg-laden germination papers were dried at 26 ± 1 °C and 70 ± 10% humidity for one week, and eggs were counted under a stereomicroscope. Eggs were hatched in 2-oz plastic soufflé cups (WebstaurantStore, Lititz, PA) filled to a height of ~1 cm of deionized water, provided fish food daily (Hikari Tropic First Bites, Petco, San Diego, CA, USA), and maintained under the same LD, temperature, and humidity conditions. Larvae were counted at 3, 4, 6, and 8 days post-hatching.

### 2.4 Locomotor activity

Spontaneous locomotor activity of female mosquitoes was recorded using a locomotor activity monitor, or actometer (Trikinetics LAM25, Waltham, MA, USA). Six to 8 day-old individual female mosquitoes that fed on one of the four test diets were placed in Pyrex glass cylindrical tubes (25 mm diameter x 125 mm length, Trikinetics, Waltham, MA, USA) immediately after feeding on one of the test diets. Tubes containing individualized mosquitoes were then placed in the activity monitor, which has three sets of infrared emitters and detectors per tube opening that bisect the approximate center of the tubes. For the duration of the experiments, mosquitoes were provided with access to 10% sucrose delivered on a soaked cotton ball placed at one end of the tube. With the exception of the sugar-fed treatment, only visibly engorged mosquitoes were used for activity experiments. Mosquitoes were maintained in the activity monitor for a duration of 5 days. Reduction in activity after blood-feeding was greatest in the first two days, as expected, since eggs are typically ready to be oviposited by three days post-feeding, so our later analysis focused on comparing those two days with the two subsequent days (*i.e.,* days 3 and 4). At the end of the experiment, egg deposition in the tubes was visually determined for each individual and, if eggs were observed, the number of eggs present was recorded. Daily locomotion was recorded as the number of beam crossings per 10-minute interval. All recordings occurred in a light-proof enclosure with its own lighting system, which consisted of a light-emitting diode (LED) light (800 Lumen, Philips, Amsterdam, The Netherlands) timed to a 12 h:12 h LD cycle aligned to the light cycle of the mosquitoes’ rearing conditions. Locomotor activity and sleep were analyzed using the *rethomics* R package as previously described (Ajayi et al., 2024, 2022; Geissmann et al., 2019; Wynne et al., 2024). Individuals that were found dead at the end of the experiments were discarded from the analysis.

### 2.5 Transcript abundance

Samples were collected by pooling 7–14 female mosquitoes (aged 7–9 days) that were visibility engorged after being provided one of the test diets, warmed to 37 □via a glass artificial feeder (Lillie Glassblowers, Smyrna, GA) during the last two hours of light during a 12 h:12 h light:dark daily light cycle (ZT 10–12). At 4 and 12 hours post□feeding, mosquitoes were decapitated using a razor blade, and heads were flash frozen in liquid nitrogen and stored at −70 □for later sample processing. Initially, expression differences were compared to blood-and sugar-fed groups, and if significant differences occurred, the Hb and noHb diets were assessed.

Total RNA was extracted from each sample using Qiagen RNeasy mini kit and RNase-free DNase (Qiagen, Hilden, Germany) following the manufacturer’s instructions. Using isolated RNA as a template, complementary DNA (cDNA) was synthesized using SuperScript III first-strand synthesis system (Thermo Fisher Scientific, Waltham, MA). Transcript levels were then quantified by determining the fluorescence of SYBR green (Thermo Fisher Scientific, Waltham, MA) of three biological replicates via quantitative real-time PCR using primers specific to each gene of interest (**Table 2**). Each reaction was run with three technical replicates. A non-template negative control was included for each primer reaction. Transcript abundance was determined by calculating the calibrated normalized relative quantities (CNRQ) following previously established guidelines (Hellemans et al., 2007; Hellemans and Vandesompele, 2011) using the housekeeping *4S ribosomal protein S7* as a reference gene, given that its expression does not significantly differ between these times of day in *Ae. aegypti* (Leming et al., 2014), which we confirmed for our laboratory colony (Lou et al., *in preparation*).

**Table 2.**
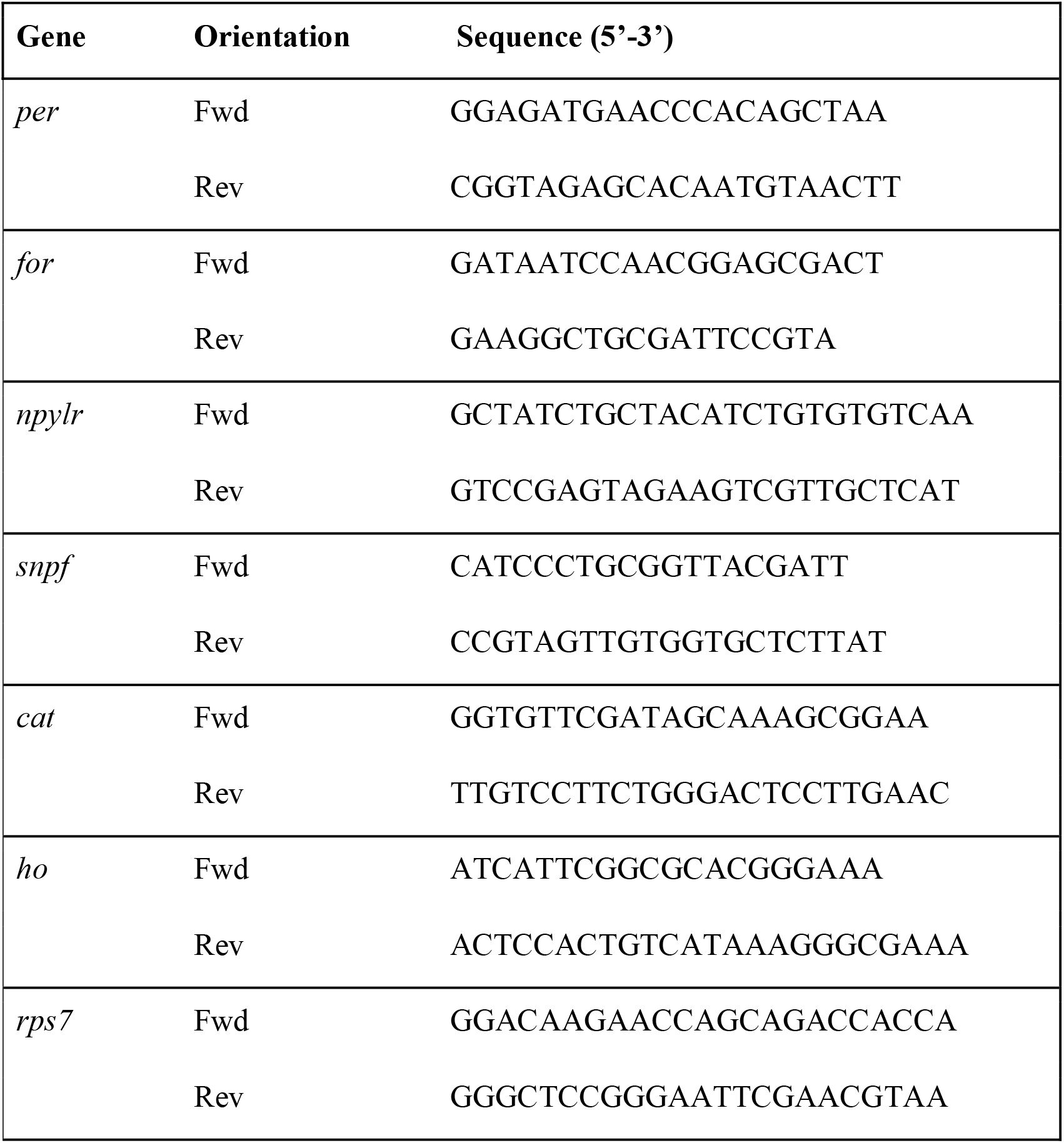
List of primers used for quantitative PCR. Primers for *npylr* are from (Liesch et al., 2013).

### 2.6. Sleep deprivation and egg production

To assess the influence of sleep deprivation on egg production, blood-fed mosquitoes were subjected to mechanical sleep disruption as previously described (Ajayi et al., 2022). First, mosquitoes were blood-fed on a human volunteer (University of Cincinnati IRB 2021-0971), placed into a mosquito cage (10 cm x 10 cm x 10 cm), and maintained under light:dark (LD) cycles of 12 h:12 h at 25 ± 1 °C and 60 ± 10% humidity. Mechanical sleep deprivation was induced using vibration stimuli (amplitude = 3 G) delivered with a Multi-Tube Vortex Mixer (Ohaus, Parsippany, NJ, USA). This vibration intensity corresponds to half the amplitude required to arouse all individuals in a previous *Drosophila* study (Shaw et al. 2000). Based on protocols adapted from another *Drosophila*-based study (Kayser et al. 2015) and studies on mosquitoes before blood feeding (Ajayi et al. 2022), two sleep deprivation regimes were applied to these diurnal mosquitoes 24 hours after receiving a blood meal: 12-hour night-time deprivation (12-NTD, sleep-deprived) and 4-hour night-time deprivation (4-NTD, negative control) with the same number of vibration periods. A sequence of vibration pulses lasting 1 min, followed by 5 min of rest between pulses, was programmed for the entire scotophase for the 12-NTD. For 4-NTD, vibration pulses lasted for 1 min followed by 1 min of rest between pulses in the first 4 h of the night (Zeitgeber time ZT12–ZT16) following the baseline day. These two conditions allow for the same number of disturbances, but allow one group 8 hours of period to sleep before dawn. After deprivation, individual females were placed into cages and allowed access to 10% sucrose and water, along with provided access to an oviposition site. Eggs were collected daily and assessed until females deposited no additional eggs for 48 hours. Fifteen mosquitoes were assessed for each condition. After one week, eggs were hatched in 2-oz plastic vials (Thornton Plastics) filled with deionized water to cover the eggs. Fish food was provided (Tertramin Goldfish flakes, Petco, Melle, Germany) and larvae were counted after 48 hours.

## 3 Results

### 3.1 Test diets

Gut dissections confirmed that mosquitoes fed on whole blood, the Hb diet, or the noHb diet ingested their meals into the midgut, with no visible accumulation in the crop (**Figure 1A**). Body mass between engorged mosquitoes did not differ signficantly across the three diets (blood-fed mass = 3.51 ± 0.14 mg; Hb-fed mass = 3.94 ± 0.39 mg; noHb-fed mass= 3.97 ± 0.27 mg, n = 15–29; Wilcoxon test, *p* > 0.1); in all cases, fed mosquitoes weighed approximately twice as much as unfed controls (Unfed mass = 1.63 ± 0.09 mg; **Figure 1B**; *p* < 0.001, *t*-test).

**Figure 1.**
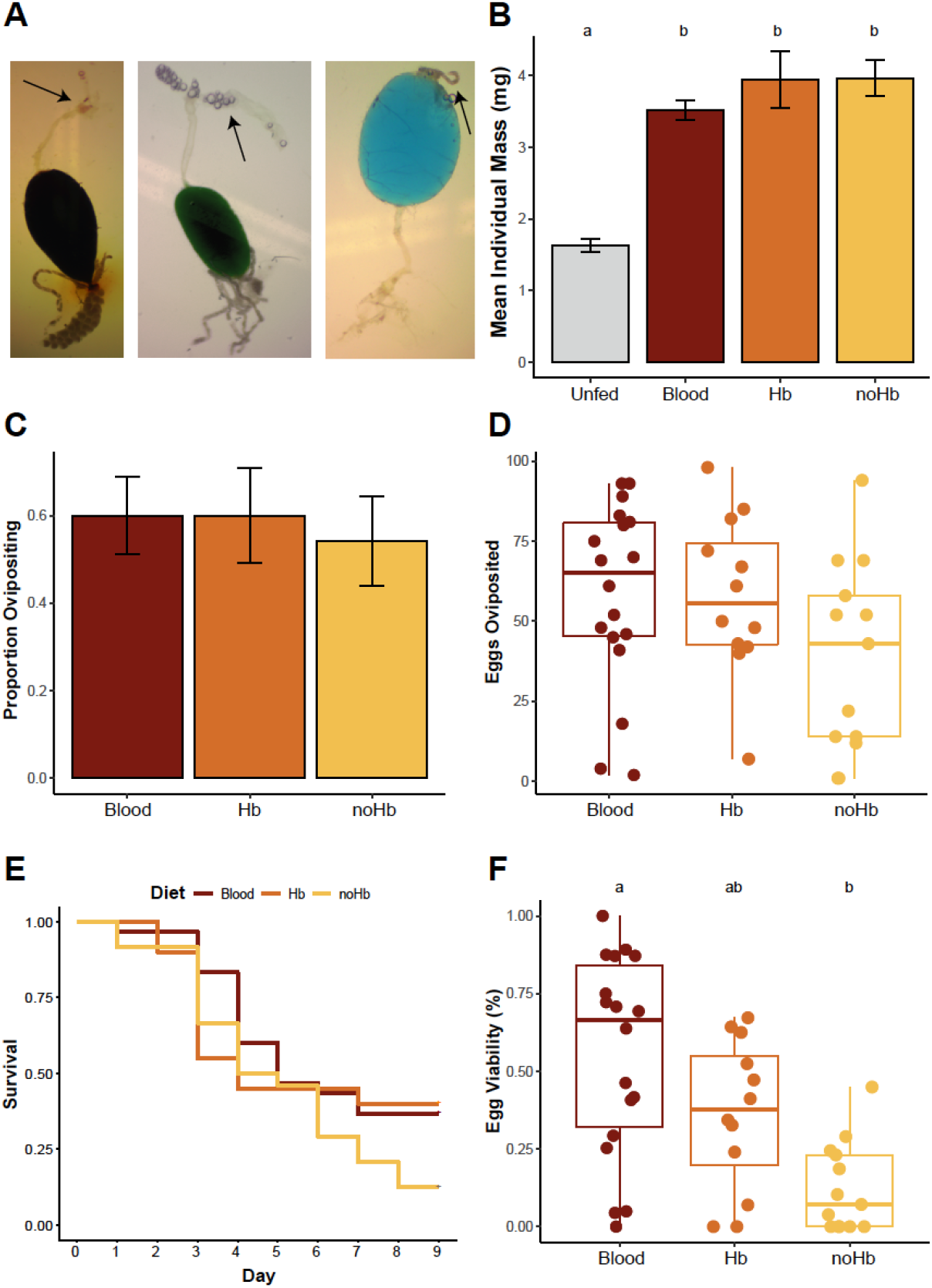
Hemoglobin is required for egg viability in *Ae. aegypti* females, although blood and artificial diets support similar egg production. **A.** Gut dissections of females fed on whole blood (*left*), Hb diet (*middle;* food dye used to enhance visibility of diet in the gut), and noHb diet (*right*; with food dye included), showing engorged midguts and crops (arrows). **B.** Mean body mass of unfed females and engorged females across diet treatments. Error bars represent the standard error of the mean. Letters denote statistical significance (*p* < 0.001) of pairwise *t*-tests performed with the Holm correction for multiple comparisons. **C.** Proportion of mosquitoes ovipositing following feeding on each of the three diets. Error bars represent the standard error of the mean. **D.** Number of eggs laid per engorged individual across diet treatments. **E.** Kaplan-Meier survival curves and log rank analysis of mosquitoes fed on each of the three diets in the oviposition assay, showing survival is not significantly different (*p* = 0.22). **F.** Egg viability (% hatched eggs by diet treatment) from the oviposition assay. Letters denote statistical significance (*p* < 0.001) of pairwise *t*-tests performed with the Holm correction for multiple comparisons.

Mosquitoes fed on blood and artificial test diets laid eggs at a similar proportion (Test of Equal Proportions, *p* > 0.1; **Figure 1C**), and there were no significant differences in the number of eggs laid between diet groups (**Figure 1D**; *p* > 0.05, multiple pairwise *t*-test with Holm *p* value adjustments), although the noHB fed females had a tendency to lay fewer eggs than blood-fed (*p* = 0.076) and Hb-fed females (*p* = 0.09). Similarly, survival rates after feeding did not differ significantly among the test diet groups (**Figure 1E**; *p* = 0.22, log-rank test). However, egg viability differed significantly by diet: mosquitoes fed the noHb diet produced fewer viable eggs compared to those fed whole blood (**Figure 1F**; *p* < 0.001, multiple pairwise *t*-test with Holm correction).

### 3.2 Locomotor activity

Feeding status and time of day significantly predicted locomotor activity (Generalized Linear Model, *p* = 0.011 for sucrose feeding and *p* = 0.006 for late evening phase). Sugar-fed females displayed the expected, stereotypical diurnal activity pattern, with a pronounced peak during the late photophase (**Figure 2A**). In contrast, blood-fed females exhibited markedly reduced activity during the first two days post-feeding, resuming minimal activity at the end of the photophase on the third day (**Figure 2A**). Females fed on either artificial diet (Hb or noHb) also showed reduced activity relative to sugar-fed controls (**Figure 2B**).

**Figure 2.**
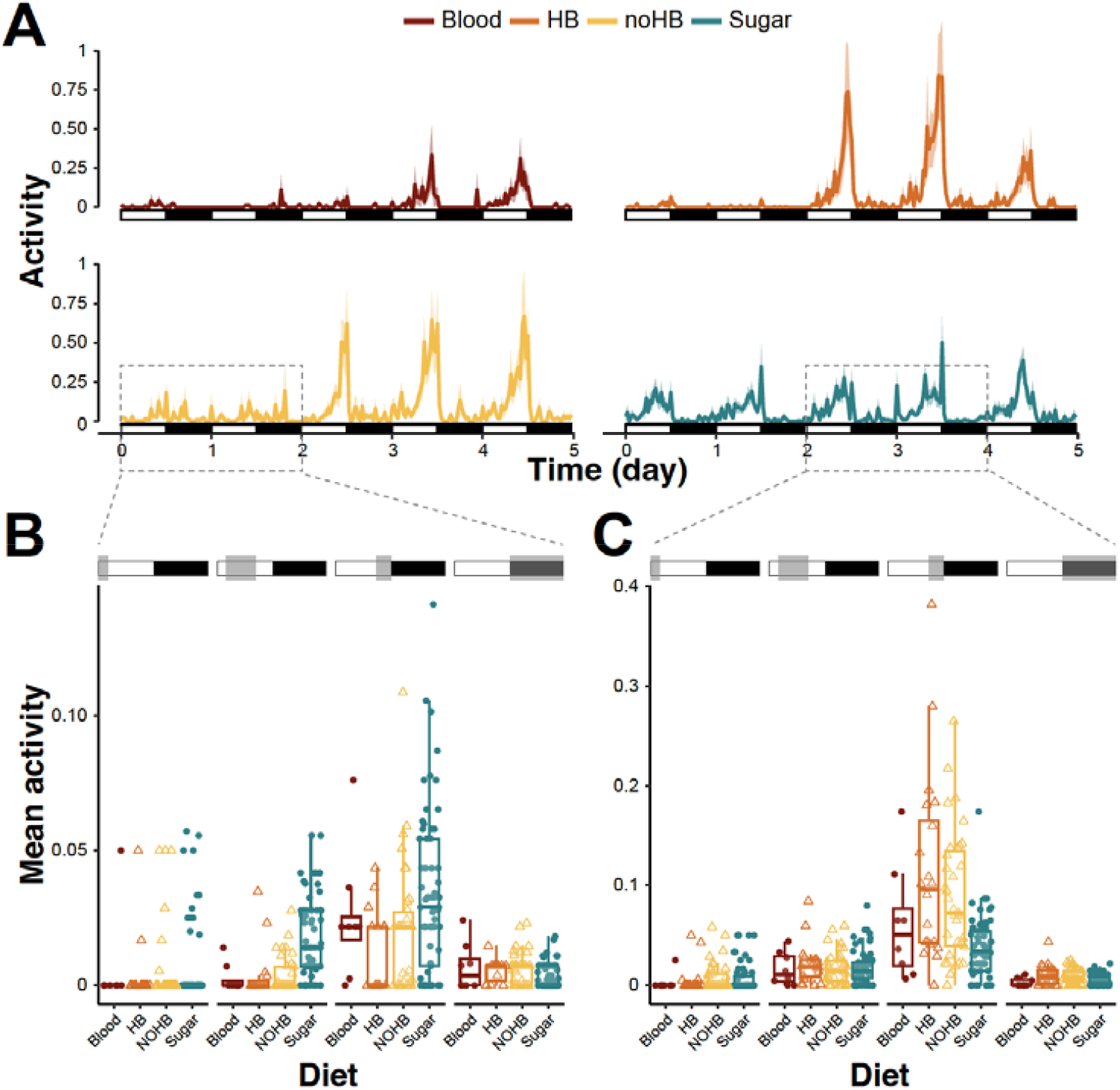
Blood and artificial diet suppress locomotor activity for two days post-feeding. **A.** Actograms of *Ae. aegypti* females fed on whole bood (n = 30), Hb diet (n = 60), no-Hb diet (n= 77), or sucrose (n = 66), monitored for five consecutive days post-feeding. Solid lines represent the mean normalized activity, and the shaded areas depict the 95% confidence intervals. **B.** Mean individual activity levels during the first two days post-feeding (highlighted by dashed box in panel A) and separated by early morning (ZT 0 - ZT 2), daytime (ZT 2 - ZT 8), end of day (ZT 8 - ZT 12), and nighttime (ZT 12 - ZT 0) phases. **C.** Mean individual activity levels during days three and four post-feeding, partitioned into the same time intervals as in panel B.. Activity levels were compared using generalized linear models with the results summarized in Tables 3 and 4.

During the stereotypical activity peak for *Ae. aegypti*, both artificial diets significantly reduced locomotor activity relative to controls (Tukey post-hoc test, *p =* 0.002 and *p* = 0.037 for the Hb and noHb diets, respectively), but the presence or absence of hemoglobin did not significantly affect activity (*p* = 0.98).

**Table 3.** Statistical summary of the diet and phase-of-day effects on mosquito activity during days one and two. (Generalized Linear Model with Tukey post hoc comparisons performed using the *emmeans* R package).

| Phase | Diet | emmean | SE | df | lower.CL | upper.CL | Letter group |
| --- | --- | --- | --- | --- | --- | --- | --- |
| daytime | Blood | 0.0026 | 0.00589 | 472 | -0.008977 | 0.01419 | ABC |
| nighttime | Sugar | 0.00337 | 0.00207 | 472 | -0.000698 | 0.00743 | A |
| daytime | NOHB | 0.00406 | 0.00286 | 472 | -0.001556 | 0.00968 | AB |
| daytime | HB | 0.0044 | 0.0043 | 472 | -0.004059 | 0.01286 | ABC |
| morning | HB | 0.00444 | 0.0043 | 472 | -0.004013 | 0.0129 | ABC |
| nighttime | HB | 0.00556 | 0.0043 | 472 | -0.002902 | 0.01401 | ABC |
| morning | NOHB | 0.0059 | 0.00286 | 472 | 0.000279 | 0.01151 | AB |
| morning | Blood | 0.00625 | 0.00589 | 472 | -0.005331 | 0.01783 | ABC |
| nighttime | NOHB | 0.00629 | 0.00286 | 472 | 0.000673 | 0.01191 | AB |
| nighttime | Blood | 0.00674 | 0.00589 | 472 | -0.004838 | 0.01832 | ABC |
| morning | Sugar | 0.007 | 0.00207 | 472 | 0.002937 | 0.01106 | AB |
| evening | HB | 0.01304 | 0.0043 | 472 | 0.004586 | 0.0215 | ABC |
| daytime | Sugar | 0.01852 | 0.00207 | 472 | 0.014456 | 0.02258 | C |
| evening | NOHB | 0.02087 | 0.00286 | 472 | 0.015257 | 0.02649 | C |
| evening | Blood | 0.02521 | 0.00589 | 472 | 0.01363 | 0.03679 | BCD |
| evening | Sugar | 0.03335 | 0.00207 | 472 | 0.029292 | 0.03742 | D |

**Table 4.** Statistical summary of the diet and phase-of-day effects on mosquito activity during days three and four. (Generalized Linear Model and Tukey post hoc comparisons performed using the *emmeans* R package).

| Phase | Diet | emmean | SE | df | lower.CL | upper.CL | Letter group |
| --- | --- | --- | --- | --- | --- | --- | --- |
| nighttime | Blood | 0.00287 | 0.01093 | 484 | -1.86E-02 | 0.0243 | AB |
| morning | Blood | 0.00313 | 0.01093 | 484 | -1.84E-02 | 0.0246 | AB |
| nighttime | Sugar | 0.00465 | 0.00393 | 484 | -3.06E-03 | 0.0124 | A |
| morning | HB | 0.00517 | 0.00691 | 484 | -8.41E-03 | 0.0188 | A |
| morning | Sugar | 0.0078 | 0.00393 | 484 | 8.66E-05 | 0.0155 | A |
| morning | NOHB | 0.0084 | 0.00523 | 484 | -1.87E-03 | 0.0187 | A |
| nighttime | NOHB | 0.00927 | 0.00523 | 484 | -9.98E-04 | 0.0195 | A |
| nighttime | HB | 0.00983 | 0.00691 | 484 | -3.76E-03 | 0.0234 | A |
| daytime | Blood | 0.01628 | 0.01093 | 484 | -5.20E-03 | 0.0378 | ABC |
| daytime | NOHB | 0.01741 | 0.00523 | 484 | 7.14E-03 | 0.0277 | AB |
| daytime | Sugar | 0.01811 | 0.00393 | 484 | 1.04E-02 | 0.0258 | A |
| daytime | HB | 0.02118 | 0.0069<br>1 | 484 | 7.60E-03 | 0.0348 | ABC |
| evening | Sugar | 0.03846 | 0.0039<br>3 | 484 | 3.07E-02 | 0.0462 | BC |
| evening | Blood | 0.06139 | 0.0109<br>3 | 484 | 3.99E-02 | 0.0829 | CD |
| evening | NOHB | 0.08841 | 0.0052<br>3 | 484 | 7.81E-02 | 0.0987 | DE |
| evening | HB | 0.11234 | 0.0069<br>1 | 484 | 9.88E-02 | 0.1259 | E |

By days three and four post-feeding, locomotor activity patterns returned to baseline, with time of day being the most important predictor of activity (Generalized Linear Model, *p* = 0.003). However, artificial diet-fed females, particularly Hb-fed individuals, displayed a rebound in activity, where the interaction between the evening phase and their diet significantly predicted this increase (Generalized Linear Model, *p* = 0.012) (**Table 3**).

By the end of the day, both Hb-fed and noHb-fed females were significantly more active than the sugar-fed controls (Tukey post-hoc test, *p* < 0.0001), and the Hb-fed females were also significantly more active than blood-fed individuals (Tukey post-hoc test, *p* = 0.009).

### 3.3 Sleep profiles

Diet and phase-of-day significantly predicted sleep amounts, with sugar diet being a significant predictor of increased sleep during the first two days post-feeding (**Figure 3A**; Generalized Linear Model, *p* = 0.020). The interaction between the end of the day (late photophase) and sugar diet was also significant (*p* = 0.006), while the interaction between noHb diet and late photophase was marginally significant (*p* = 0.057).

**Figure 3.**
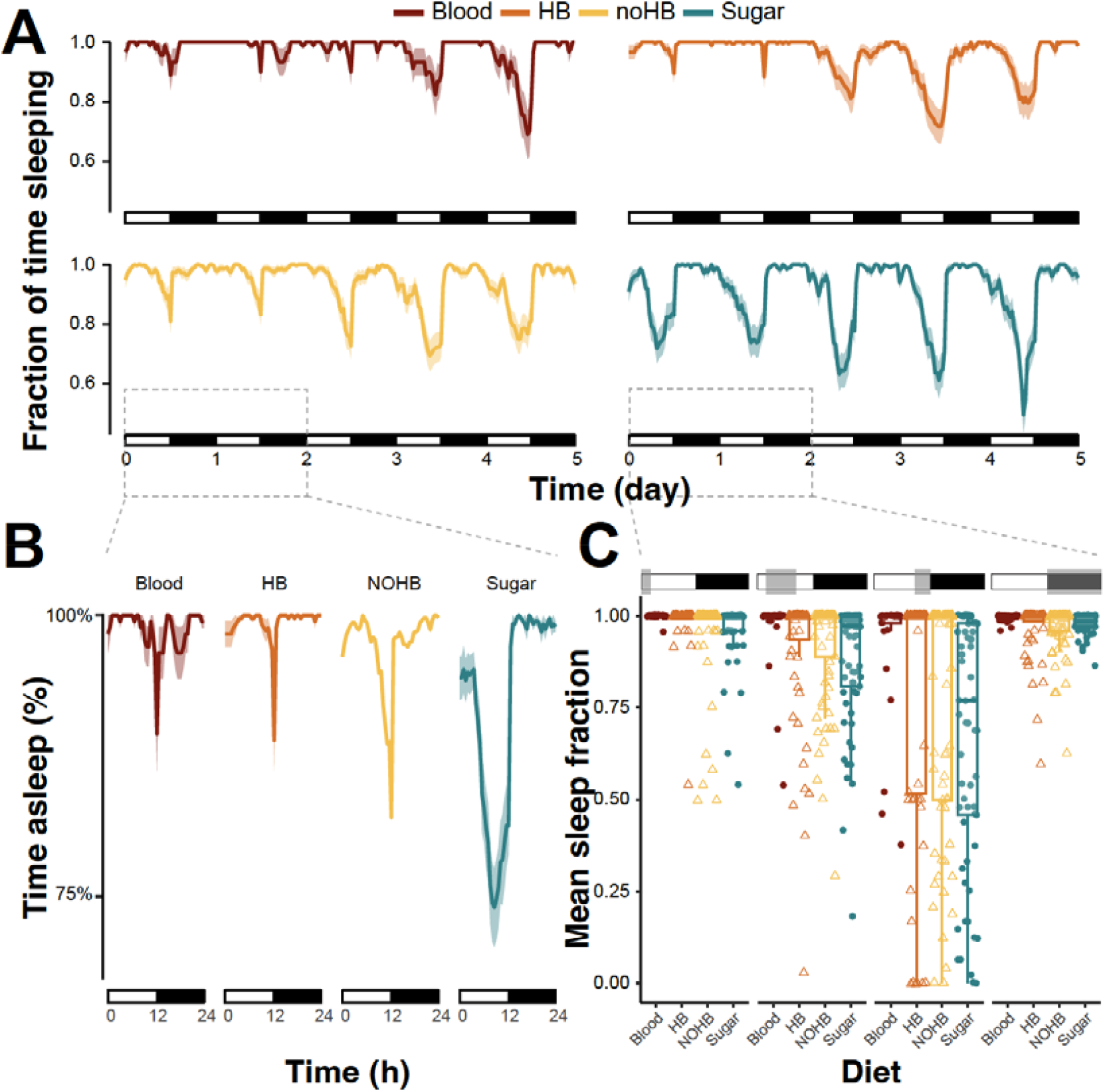
Hemoglobin-containing diets and blood increase post-feeding sleep propensity. **A.** Sleep profiles of *Ae. aegypti* females fed on whole blood (n = 30), Hb diet (n = 60), noHb diet (n= 77), or sucrose (n = 66), monitored for five consecutive days post-feeding. Solid lines represent the mean fraction of time spent in a sleep-like state, and the shaded areas depict the 95% confidence intervals. **B.** Mean sleep profiles during the first two days post-feeding (highlighted by the dashed box in panel A) for mosquitoes fed either artificial diet, visualized over a 24-hour cycle. **C.** Mean fraction of time spent sleeping per individual, separated into early morning (ZT 0-2), daytime (ZT 2-8), end of day (ZT 8-12), and nighttime (ZT 12-0). Statistical results are summarized in Table 5.

At the end of the day (photophase), Hb-fed mosquitoes slept significantly more than sugar-fed controls (**Figure 3C**; Tukey post hoc test, *p* = 0.042), but their sleep duration did not differ significantly from that of the blood-fed mosquitoes (Tukey post hoc test, *p* = 0.065).

**Table 5.** Statistical summary of the diet and phase-of-day effects on mosquito sleep during days one and two. (Generalized Linear Model and Tukey post hoc comparisons performed using the *emmeans* R package).

| Phase | Diet | emm<br>ean | SE | df | lower.CL | upper.C<br>L | Letter<br>group |
| --- | --- | --- | --- | --- | --- | --- | --- |
| evening | Sugar | 0.676 | 0.0227 | 916 | 0.632 | 0.721 | A |
| evening | NOHB | 0.775 | 0.021 | 916 | 0.733 | 0.816 | AB |
| evening | HB | 0.791 | 0.0238 | 916 | 0.744 | 0.837 | BC |
| daytime | Sugar | 0.874 | 0.0227 | 916 | 0.829 | 0.918 | BCD |
| daytime | HB | 0.91 | 0.0238 | 916 | 0.863 | 0.956 | DE |
| daytime | NOHB | 0.921 | 0.021 | 916 | 0.88 | 0.962 | DE |
| evening | Blood | 0.928 | 0.0336 | 916 | 0.863 | 0.994 | CDE |
| morning | NOHB | 0.961 | 0.021 | 916 | 0.92 | 1.002 | DE |
| morning | Sugar | 0.963 | 0.0227 | 916 | 0.918 | 1.007 | DE |
| nighttime | HB | 0.966 | 0.0238 | 916 | 0.919 | 1.012 | DE |
| nighttime | NOHB | 0.967 | 0.021 | 916 | 0.926 | 1.008 | DE |
| daytime | Blood | 0.968 | 0.0336 | 916 | 0.902 | 1.034 | DE |
| nighttime | Sugar | 0.979 | 0.0227 | 916 | 0.935 | 1.024 | DE |
| morning | HB | 0.988 | 0.0238 | 916 | 0.941 | 1.034 | E |
| nighttime | Blood | 0.994 | 0.0336 | 916 | 0.928 | 1.06 | DE |
| morning | Blood | 0.999 | 0.0336 | 916 | 0.933 | 1.065 | DE |

In contrast, noHb-fed mosquitoes slept significantly less than blood-fed individuals (Tukey post hoc test, *p* = 0.010) but not significantly more than the sugar-fed individuals (*p* = 0.104). No significant differences were observed between the sleep amounts of the Hb-fed and noHb-fed groups (*p* > 0.05).

### 3.4 Mosquito head transcript abundance

Expression of the core circadian gene *period (per)* in mosquito heads was significantly reduced 4 hours after feeding on either whole blood or the Hb diet, relative to sugar-fed controls (**Figure 4A**; *p* < 0.05, *t*-test). No significant differences in *per* expression were observed at 12 hours post-feeding (**Figure 4B**). In noHB-fed females, *per* expression did not differ significantly from either sugar-fed nor Hb-fed groups.

**Figure 4.**
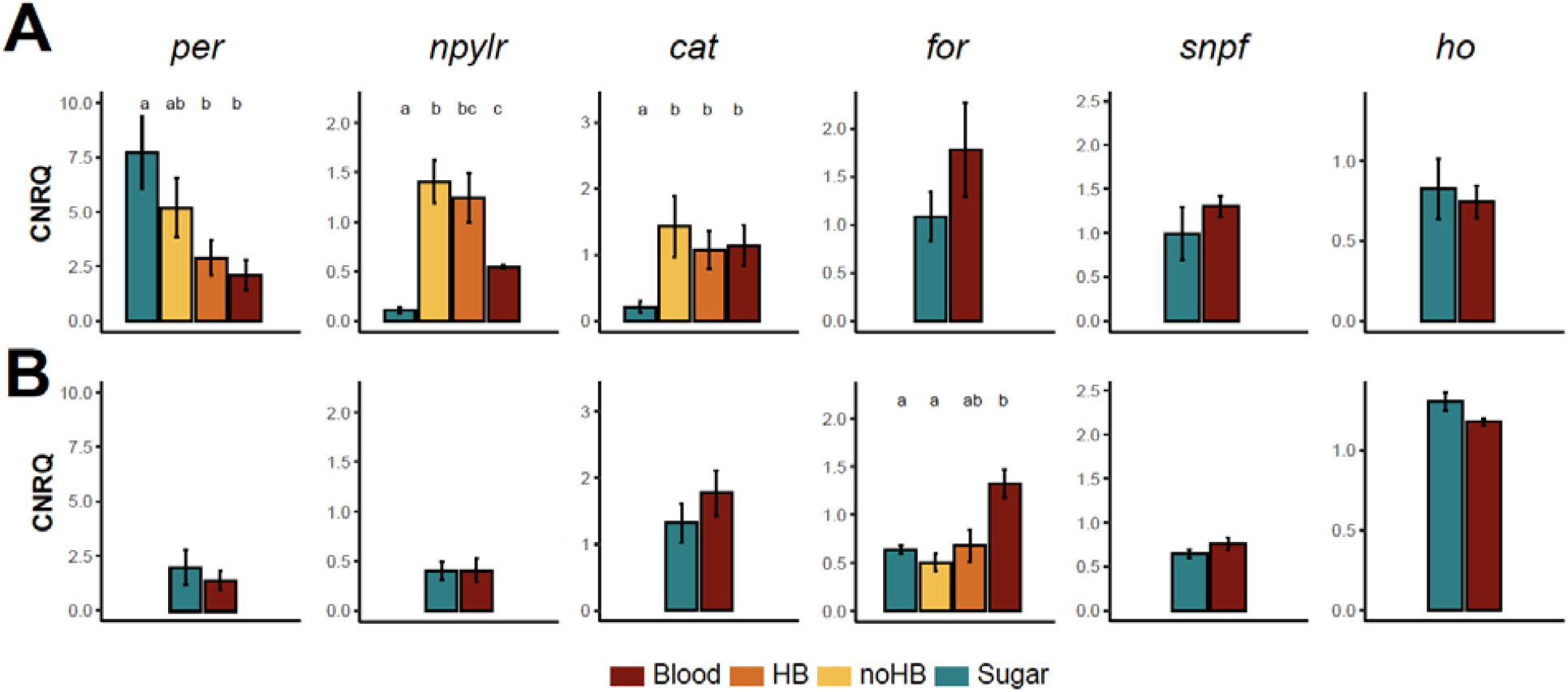
Transcript abundance of select clock, host seeking, and redox processing is affected by the diet. **A.** Transcript levels of (from *left* to *right*) *period, neuropeptide Y-like receptor I, catalase, foraging, short neuropeptide*, and *heme oxygenase* in mosquito heads, 4 hours after feeding on sugar (blue), no-Hb (dark orange), Hb (orange), or blood (red) diets. Expression levels are shown as calibrated normalized relative quantities (CNRQ) normalized to the mean of all samples per gene. Transcript levels in mosquitoes fed on Hb or noHb diets were quantified only when expression levels differed between sugar- and blood-fed mosquitoes (*p* < 0.05, *t*-test performed on log2(CNRQ) values). **B.** Transcript abundance quantified for the same genes as in A at 12 hours post-feeding. Error bars represent the standard error of the mean. Letters denote statistical significance (*p* < 0.05) of pairwise *t*-tests performed with the Holm correction for multiple comparisons.

At 4 hours post-feeding, transcript abundance of *Neuropeptide Y-like receptor (npylr)* was significantly elevated in all three non-sugar-fed groups (whole blood, Hb, and noHb) compared to sugar-fed controls (**Figure 4A**; *p* < 0.05, *t*-test), with no differences observed at 12 hours post-meal.

Similarly, the expression of the antioxidant gene *catalase (cat)* was significantly upregulated 4 hours post feeding in all non-sugar-fed groups (whole blood, noHb, and Hb) relative to sugar-fed mosquitoes (**Figure 4A**; *p* < 0.05, *t*-test), but not 12 hours post feeding (**Figure 4B**).

In contrast, *foraging (for)* transcript levels were significantly elevated in blood-fed mosquitoes 12 hours post-feeding compared to sugar-fed and noHB-fed groups (**Figure 4B**; *p* < 0.05, *t*-test). Expression of *for in* noHb-fed mosquitoes was not significantly different from sugar-fed controls while Hb-fed mosquitoes exhibited intermediate *for* expression, but differences were not statistically significant.

No significant differences in the expression of *short neuropeptide F (snpf)* or *heme oxygenase (ho)* were detected at either time point post-feeding (**Figure 4A,B**; *p* > 0.05, *t*-test).

### 3.5 Sleep deprivation

As previously noted, there is a reduction in activity and increased sleep post-blood feeding associated with the ingestion of hemoglobin and blood. We validated these patterns following feeding on a live human host, which showed the same activity reduction and increased sleep during the first two days after a bloodmeal (**Figure 5A, B**). During the nighttime 24-h post blood-feeding, mosquitoes were subjected to sleep deprivation as previously described (Ajayi et al., 2022), but this did not impact subsequent egg production - either total or daily output (**Figure 5C**) of females. There was no significant difference in the number of mosquitoes that produced eggs between the groups, ranging from 90-100% of mosquitoes yielded eggs for each treatment. Lastly, the viability of the eggs did not vary between groups, with 80-82% viability for the egg among all three treatments (*p* > 0.80). This observation suggests that a single night of sleep deprivation is unlikely to reduce either the timing of egg deposition, the number of egg produced, or egg viability.

**Figure 5.**
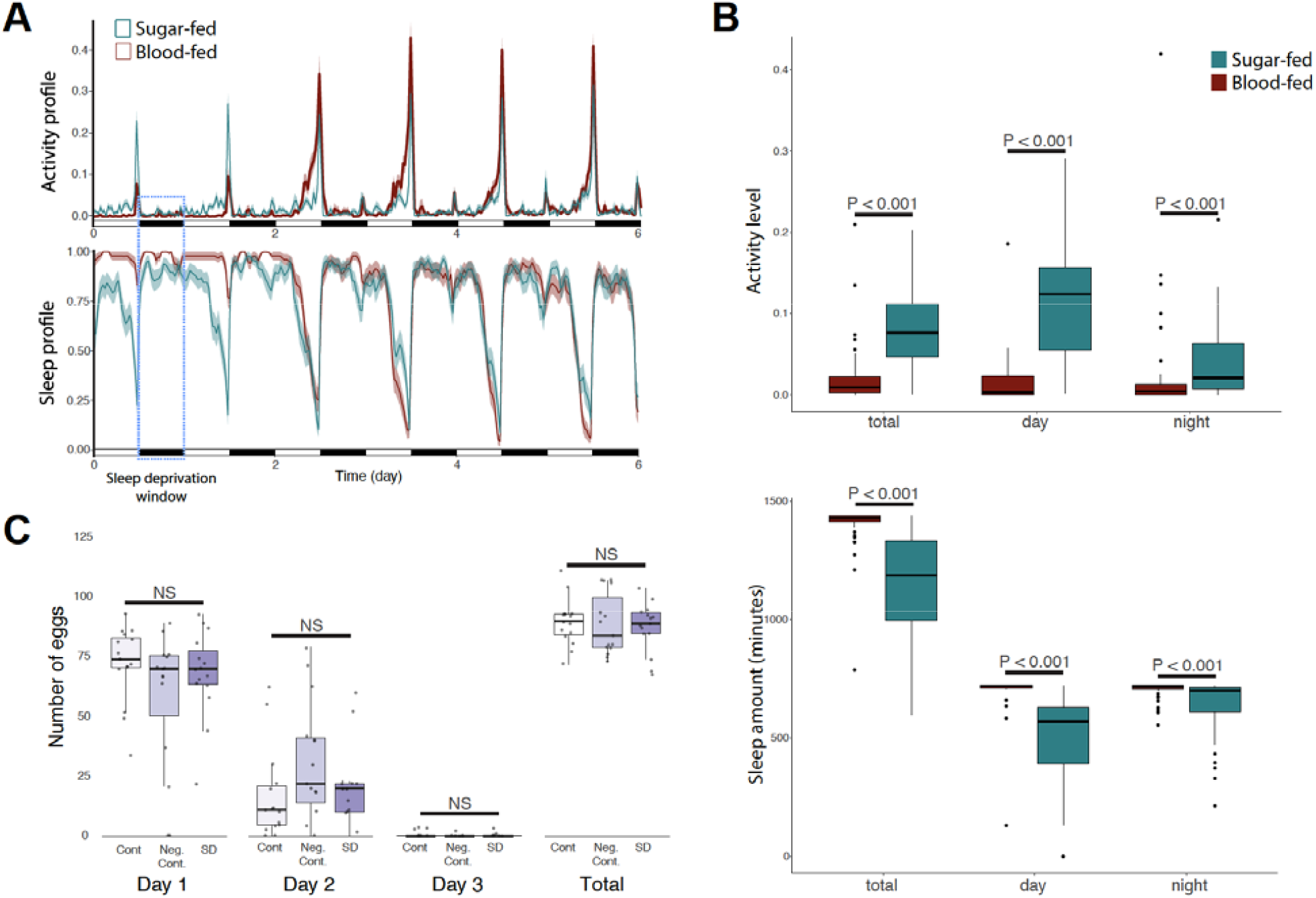
Sleep levels following a bloomeal and egg production in relation to sleep deprivation for *Aedes aegypti*. **A.** Baseline activity and sleep rhythm of *Aedes aegypti* after a bloodmeal. The y-axis represents the mean beam crosses in an activity monitor made by all the mosquitoes or the proportion of time spent sleeping (defined as inactive periods of 120 min), averaged for each mosquito within a 30 min time window (N=25-30). **B.** Amount of activity and sleep in total and during the day and night in the first day after a bloodmeal (N=25-30) compared to sugar-fed individuals. Significance based on a Wilcoxon rank sum test with continuity correction. **C.** Egg production following sleep deprivation (denoted by blue box in A), N =15 per treatment. Control (Cont) = no mechanical vibration, Negative Control (Neg Cont), mechanical vibration limited to a 4 hour period to allow mosquitoes to sleep the remaining 8 hours of the night, and sleep-deprived (SD) mechanical vibration during the entire night. No differences were noted in the number of females that deposited eggs and egg viability (P > 0.6 in all cases). Statistical differences were assessed with a general linear model for egg number differences (Gaussian).

## 4. Discussion

In this study, we took advantage of an artificial diet originally proposed as an alternate source of proteins to maintain mosquitoes in the laboratory, bypassing the need for animal blood (Gonzales et al., 2018). Here, we used it as a means to experimentally manipulate the presence or absence of hemoglobin in the meal. Our results show that the lack of heme has no effect on mosquitoes’ survival, propensity to feed, or the amount of fluid ingested when compared to bovine blood provided via an artificial membrane feeder. However, the absence of hemoglobin led to a slight reduction in the number of eggs produced and a significant decrease in their viability (**Figure 1**). Given that a certain amount of iron is required to produce viable eggs (~7% of the ingested heme) (Zhou et al., 2007), the observed phenotype could result from an incomplete nutritional input, rather than being due to redox-dependent processes. Although not tested here, it is possible that these no-Hb-fed individuals would be seeking a secondary meal to compensate for the insufficient heme acquisition and behave like partially-fed mosquitoes (Klowden and Lea, 1978).

Similar to blood, artificial diets reduced locomotor activity, particularly during the first two days post-meal (**Figure 2**). During the following two days, Hb and no-Hb diet-fed mosquitoes showed a rebound in activity that was also observed by others in response to a sheep blood meal (Dong et al., 2025) and hypothesized as an increased search for oviposition sites by females with mature eggs. Interestingly, we did not observe this activity rebound when feeding mosquitoes with bovine blood (**Figure 2**), but only when females were fed on the arms of human volunteers (**Figure 5**). These observations suggest a potential role of the blood composition (i.e., potentially its nutritional value) in leading to this activity rebound, suggesting that artificial feeding, even though sufficient for egg production, may mask specific behavioral shifts post-blood feeding if a live host is used.

When analyzing sleep, a long-overlooked aspect of mosquito activity (Ajayi et al., 2023), we found that heme is responsible to a large extent for the increase in sleep propensity observed in the first two days post blood-feeding, as similar sleep amounts were recapitulated in the Hb-fed individuals, but not in the no-Hb individuals, which slept comparably to their sugar-fed counterparts (**Figure 3**). This suggests that post-feeding increases sleep-like states could require the ingestion of heme, which acts as a critical component of the bloodmeal in producing eggs (Zhou et al., 2007). This lack of mosquitoes in a sleep-like state could increase the potential for a secondary blood feeding, which has been observed when mosquitoes fail to locate a stable resting habitat after a bloodmeal (Holmes et al., 2025) or ingest an insufficient initial bloodmeal (Duvall, 2019; Scott et al., 1993).

At the transcriptional level, these behavioral effects were correlated with rapid (within 4 hours) changes in *period*, a core clock gene, and *catalase,* an essential gene for oxidative stress protection (DeJong et al., 2007; Oliveira et al., 2017). Genes directly implicated in regulating *Ae. aegypti*’s host-seeking behavior and locomotor activity (*npylr* and *for*) were also differentially expressed at 4 and 12 hours post-bloodmeal, respectively. Interestingly, in *Drosophila melanogaster,* where the *foraging* gene was first described, higher *foraging gene* expression is associated with longer foraging path lengths in larvae (Anreiter and Sokolowski, 2019). Here, the increased expression in blood and Hb-fed mosquitoes at 12 hours post-meal may suggest its potential implication in the upcoming rebound in locomotor activity and oviposition site seeking behavior, rather than its involvement in the reduction of activity that follows the meal (**Figure 4B**).

The effects on *per, catalase*, and *npylr* were largely recapitulated (or amplified in the case of *npylr*) by the artificial diets, although the no-Hb diet produced an intermediate phenotype for *per* (**Figure 4A**). Similarly, the no-Hb diet results in expression levels similar to those of sugar-fed mosquitoes, while the Hb-fed mosquitoes produced an intermediate expression profile between those of sugar and blood-fed mosquitoes (**Figure 4B**). *per* encodes a core clock component, and its expression patterns are known to correlate with behavioral rhythms in *Ae. aegypti* and other insects (Leming et al., 2014). Downregulation of *per* in blood- and Hb-fed females suggests a molecular mechanism through which dietary heme suppresses behavioral rhythms. Similarly, *for* encodes a cGMP-dependent protein kinase that modulates foraging and locomotor activity across insect species (Anreiter and Sokolowski, 2019); its suppression further supports the observed reduction in activity. In contrast, *npylr1*, a receptor previously implicated in host-seeking suppression (Duvall et al., 2019; Liesch et al., 2013), did not show significant differences between the Hb and no-Hb treatments, suggesting it may not be regulated by dietary heme but responds to other nutrient intake (e.g., proteins) instead.

Altogether, our results support the hypothesis that dietary heme contributes explicitly to the reduction in postprandial activity and increased sleep propensity via selective changes in the expression of genes involved in circadian rhythms, host-seeking, and oxidative stress management. This behavioral suppression may reflect an adaptive mechanism to reduce host-seeking after a successful blood meal, thus conserving energy for digestion and egg development and potentially reducing exposure to predators or further host encounters (Klowden and Lea, 1979, 1978; Takken et al., 2001). This period of suppressed activity requires the post-blood females to find a stable and humid location to rest, where subpar conditions can result in an early return of activity and a secondary feeding (Holmes et al., 2025). Our results suggest that this suppression may be driven at least in part by the consumption of sufficient dietary heme, and not simply by abdominal distention, protein content, or egg development.

To test whether the sleep increase induced by blood ingestion is critical for optimal reproductive output, we exposed blood-fed females to sleep deprivation post-blood feeding. In these experiments, we did not observe a decrease in egg production and viability (**Figure 5**). This suggests that the increased sleep propensity may be an adaptive mechanism to keep mosquitoes out of harm’s way (by reducing their risks of encountering a predator or a defensive host at a time when the additional blood payload impairs their mobility) rather than a response to a physiological need to process blood nutrients for egg production. It is worth noting, however, that a more extended period of reduced sleep (24-36 hours) could have a more substantial impact on digestion and egg output and warrants further study. Such experiments are challenging given the increased mortality observed in prolonged sleep reduction assays (Ajayi et al., 2022). Alternatively, transfer of specific nutrients (lipid, proteins, or carbohydrate) could have been impacted, and the nutritional composition of eggs, as well as the fate of artificial diet-fed progeny, represents another target for future studies.

In conclusion, our findings support a model in which dietary heme, delivered via hemoglobin, plays a key role in modulating behavior and gene expression in *Ae. aegypti* females following a blood meal. These results highlight a previously underappreciated function of heme in coordinating the suppression of host-seeking and activation of antioxidant defenses. While our results point to a central role of dietary heme in regulating mosquito behavior and gene expression, we also observed some distinctions between blood- and Hb-fed mosquitoes. This indicates that other differences between these fluids, such as lipids, micronutrients, or signaling molecules composition, could also influence post-feeding physiology and behavior. Given the central importance of blood-feeding in the transmission of vector-borne diseases, a better understanding of the physiological and molecular responses triggered by dietary components such as heme may inform novel strategies for mosquito control.

## Author Contributions

Conceptualization, D.F.E. and C.V.; methodology, D.F.E, K.C., and C.V.; formal analysis, D.F.E., J.B.B., and C.V.; investigation, D.F.E., K.C., O.E., A.S., O.M.A., M.V., and C.V.; writing—original draft preparation, D.F.E.; writing—review and editing, D.F.E., J.B.B., and C.V.; visualization, D.F.E., O.M.A., J.B.B., and C.V.; supervision, C.V.; project administration, C.V.

## Acknowledgements

We thank the Fralin Life Sciences Institute and the Department of Biochemistry for material support and access to shared equipment used in this study, D. Dougharty for his assistance with mosquito care, and C. Lahondère for helpful suggestions.

## References

Ajayi, O.M., Eilerts, D.F., Bailey, S.T., Vinauger, C., Benoit, J.B., 2020. Do Mosquitoes Sleep? Trends Parasitol. 36, 888–897.

Ajayi, O.M., Marlman, J.M., Gleitz, L.A., Smith, E.S., Piller, B.D., Krupa, J.A., Vinauger, C., Benoit, J.B., 2022. Behavioral and postural analyses establish sleep-like states for mosquitoes that can impact host landing and blood feeding. J. Exp. Biol. 225. 10.1242/jeb.244032

Ajayi, O.M., Susanto, E.E., Wang, L., Kennedy, J., Ledezma, A., Harris, A., Smith, E.S., Chakraborty, S., Wynne, N.E., Sylla, M., Akorli, J., Otoo, S., Rose, N.H., Vinauger, C., Benoit, J.B., 2024. Intra-species quantification reveals differences in activity and sleep levels in the yellow fever mosquito, Aedes aegypti. Medical and Veterinary Entomology. 10.1111/mve.12747

Ajayi, O.M., Wynne, N.E., Chen, S.-C., Vinauger, C., Benoit, J.B., 2023. Sleep: An essential and understudied process in the biology of blood-feeding arthropods. Integr. Comp. Biol. 10.1093/icb/icad097

Anreiter, I., Sokolowski, M.B., 2019. The foraging gene and its behavioral effects: Pleiotropy and plasticity. Annu. Rev. Genet. 53, 373–392.

Barredo, E., DeGennaro, M., 2020. Not just from blood: mosquito nutrient acquisition from nectar sources. Trends Parasitol. 36, 473–484.

Benoit, J.B., Denlinger, D.L., 2017. Bugs battle stress from hot blood. Elife 6, e33035.

Benoit, J.B., Lopez-Martinez, G., Phillips, Z.P., Patrick, K.R., Denlinger, D.L., 2010. Heat shock proteins contribute to mosquito dehydration tolerance. J. Insect Physiol. 56, 151–156.

Briegel, H., 1985. Mosquito reproduction: Incomplete utilization of the blood meal protein for oögenesis. J. Insect Physiol. 31, 15–21.

Chaitanya, R.K., Shashank, K., Sridevi, P., 2016. Oxidative Stress in Invertebrate Systems, in: Ahmad, R. (Ed.), Free Radicals and Diseases. InTech.

de Lima-Camara, T.N., 2010. Activity patterns of Aedes aegypti and Aedes albopictus (diptera: culicidae) under natural and artificial conditions. Oecologia Australis 14, 737–736.

DeJong, R.J., Miller, L.M., Molina-Cruz, A., Gupta, L., Kumar, S., Barillas-Mury, C., 2007. Reactive oxygen species detoxification by catalase is a major determinant of fecundity in the mosquito Anopheles gambiae. Proc. Natl. Acad. Sci. U. S. A. 104, 2121–2126.

Dong, L., Bradford, E.F., Barnett, J.M., Duvall, L.B., 2025. Post-biting behavioral reprogramming underlies reproductive efficiency in Aedes aegypti mosquitoes. Neuroscience.

Duvall, L.B., 2019. Mosquito host-seeking regulation: targets for behavioral control. Trends Parasitol. 35, 704–714.

Duvall, L.B., Ramos-Espiritu, L., Barsoum, K.E., Glickman, J.F., Vosshall, L.B., 2019. Small-molecule agonists of Ae. aegypti neuropeptide Y receptor block mosquito biting. Cell 176, 687-701.e5.

Eilerts, D.F., VanderGiessen, M., Bose, E.A., Broxton, K., Vinauger, C., 2018. Odor-specific daily rhythms in the olfactory sensitivity and behavior of Aedes aegypti mosquitoes. Insects 9, 147.

Evans, O., Caragata, E.P., McMeniman, C.J., Woolfit, M., Green, D.C., Williams, C.R., Franklin, C.E., O’Neill, S.L., McGraw, E.A., 2009. Increased locomotor activity and metabolism of Aedes aegypti infected with a life-shortening strain of Wolbachia pipientis. J. Exp. Biol. 212, 1436–1441.

Foster, W.A., 1995. Mosquito sugar feeding and reproductive energetics. Annu. Rev. Entomol. 40, 443–474.

Geissmann, Q., Garcia Rodriguez, L., Beckwith, E.J., Gilestro, G.F., 2019. Rethomics: An R framework to analyse high-throughput behavioural data. PLoS One 14, e0209331.

Gentile, C., Rivas, G.B. da S., Lima, J.B.P., Bruno, R.V., Peixoto, A.A., 2013. Circadian clock of Aedes aegypti: effects of blood-feeding, insemination and RNA interference. Mem. Inst. Oswaldo Cruz 108 Suppl 1, 80–87.

Gonzales, K.K., Rodriguez, S.D., Chung, H.-N., Kowalski, M., Vulcan, J., Moore, E.L., Li, Y., Willette, S.M., Kandel, Y., Van Voorhies, W.A., Holguin, F.O., Hanley, K.A., Hansen, I.A., 2018. The effect of skitosnack, an artificial blood meal replacement, on Aedes aegypti life history traits and gut microbiota. Sci. Rep. 8, 11023.

Greenberg, J., 1951. Some nutritional requirements of adult mosquitoes (Aedes aegypti) for oviposition. J. Nutr. 43, 27–35.

Hardeland, R., Coto-Montes, A., Poeggeler, B., 2003. Circadian rhythms, oxidative stress, and antioxidative defense mechanisms. Chronobiol. Int. 20, 921–962.

Hellemans, J., Mortier, G., De Paepe, A., Speleman, F., Vandesompele, J., 2007. qBase relative quantification framework and software for management and automated analysis of real-time quantitative PCR data. Genome Biol. 8, R19.

Hellemans, J., Vandesompele, J., 2011. qPCR data analysis-unlocking the secret to successful results. PCR troubleshooting and optimization: the essential guide 1, 13.

Holmes, C.J., Chakraborty, S., Ajayi, O.M., Uhran, M.R., Frigard, R., Stacey, C.L., Susanto, E.E., Chen, S.-C., Rasgon, J.L., DeGennaro, M., Xiao, Y., Benoit, J.B., 2025. Multiple blood feeding bouts in mosquitoes allow for prolonged survival and are predicted to increase viral transmission during dry periods. iScience 28, 111760.

Hwangbo, D.-S., Kwon, Y.-J., Iwanaszko, M., Jiang, P., Abbasi, L., Wright, N., Alli, S., Hutchison, A.L., Dinner, A.R., Braun, R.I., Allada, R., 2023. Dietary restriction impacts peripheral circadian clock output important for longevity in Drosophila. 10.7554/elife.86191.1

Kawada, H., Takemura, S.-Y., Arikawa, K., Takagi, M., 2005. Comparative study on nocturnal behavior of Aedes aegypti and Aedes albopictus. J. Med. Entomol. 42, 312–318.

Kayser, M.S., Mainwaring, B., Yue, Z., Sehgal, A., 2015. Sleep deprivation suppresses aggression in Drosophila. Elife 4, e07643.

Klowden, M.J., Lea, A., 1979. Inhibition during of host-seeking in Aedes oocyte maturation. J. Insect Physiol. 25, 231–235.

Klowden, M.J., Lea, A.O., 1978. Blood meal size as a factor affecting continued host-seeking by Aedes aegypti (L.). Am. J. Trop. Med. Hyg. 27, 827–831.

Krishnan, N., Davis, A.J., Giebultowicz, J.M., 2008. Circadian regulation of response to oxidative stress in Drosophila melanogaster. Biochem. Biophys. Res. Commun. 374, 299–303.

Lahondère, C., Lazzari, C.R., 2012. Mosquitoes cool down during blood feeding to avoid overheating. Curr. Biol. 22, 40–45.

Leming, M.T., Rund, S.S.C., Behura, S.K., Duffield, G.E., O’Tousa, J.E., 2014. A database of circadian and diel rhythmic gene expression in the yellow fever mosquito Aedes aegypti. BMC Genomics 15, 1128.

Liesch, J., Bellani, L.L., Vosshall, L.B., 2013. Functional and genetic characterization of neuropeptide Y-like receptors in Aedes aegypti. PLoS Negl. Trop. Dis. 7, e2486.

Lima-Camara, T.N., Bruno, R.V., Luz, P.M., Castro, M.G., Lourenço-de-Oliveira, R., Sorgine, M.H.F., Peixoto, A.A., 2011. Dengue infection increases the locomotor activity of Aedes aegypti females. PLoS One 6, e17690.

Lima-Camara, T.N., Lima, J.B.P., Bruno, R.V., Peixoto, A.A., 2014. Effects of insemination and blood-feeding on locomotor activity of Aedes albopictus and Aedes aegypti (Diptera: Culicidae) females under laboratory conditions. Parasit. Vectors 7, 304.

Lou, L., Chandrasegaran, K., Devilliers, J., Kobiowu, A., Compton, A. Wynne, N.E., Bisese, A., Applebach, E., Sapkota, S., Rust, R., Luff, S., Eilerts, D.F., Evans, O., Benoit, J.B., Tu, Z.J., Lahondère, C., Vinauger, C. Temporal synchrony between human odor rhythms and mosquito olfactory preference. in preparation.

Luz, P.M., Lima-Camara, T.N., Bruno, R.V., Castro, M.G. de, Sorgine, M.H.F., Lourenço-de-Oliveira, R., Peixoto, A.A., 2011. Potential impact of a presumed increase in the biting activity of dengue-virus-infected Aedes aegypti (Diptera: Culicidae) females on virus transmission dynamics. Mem. Inst. Oswaldo Cruz 106, 755–758.

Mandilaras, K., Missirlis, F., 2012. Genes for iron metabolism influence circadian rhythms in Drosophila melanogaster. Metallomics 4, 928–936.

Oliveira, J.H.M., Talyuli, O.A.C., Goncalves, R.L.S., Paiva-Silva, G.O., Sorgine, M.H.F., Alvarenga, P.H., Oliveira, P.L., 2017. Catalase protects Aedes aegypti from oxidative stress and increases midgut infection prevalence of Dengue but not Zika. PLoS Negl. Trop. Dis. 11, e0005525.

Oliver, S.V., Brooke, B.D., 2016. The role of oxidative stress in the longevity and insecticide resistance phenotype of the major malaria vectors Anopheles arabiensis and Anopheles funestus. PLoS One 11, e0151049.

Peach, D.A.H., Gries, G., 2020. Mosquito phytophagy – sources exploited, ecological function, and evolutionary transition to haematophagy. Entomol. Exp. Appl. 168, 120–136.

Pereira, L.O.R., Oliveira, P.L., Almeida, I.C., Paiva-Silva, G.O., 2007. Biglutaminyl-biliverdin IX alpha as a heme degradation product in the dengue fever insect-vector Aedes aegypti. Biochemistry 46, 6822–6829.

Rivera-Pérez, C., Clifton, M.E., Noriega, F.G., 2017. How micronutrients influence the physiology of mosquitoes. Curr. Opin. Insect Sci. 23, 112–117.

Rudisill, S.S., Martin, B.R., Mankowski, K.M., Tessier, C.R., 2019. Iron deficiency reduces synapse formation in the Drosophila clock circuit. Biol. Trace Elem. Res. 189, 241–250.

Saeaue, L., Morales, N.P., Komalamisra, N., Morales Vargas, R.E., 2011. Antioxidative systems defense against oxidative stress induced by blood meal in Aedes aegypti. Southeast Asian J. Trop. Med. Public Health 42, 542–549.

Scaraffia, P.Y., Isoe, J., Murillo, A., Wells, M.A., 2005. Ammonia metabolism in Aedes aegypti. Insect Biochem. Mol. Biol. 35, 491–503.

Scaraffia, P.Y., Tan, G., Isoe, J., Wysocki, V.H., Wells, M.A., Miesfeld, R.L., 2008. Discovery of an alternate metabolic pathway for urea synthesis in adult Aedes aegypti mosquitoes. Proc. Natl. Acad. Sci. U. S. A. 105, 518–523.

Scaraffia, P.Y., Wells, M.A., 2003. Proline can be utilized as an energy substrate during flight of Aedes aegypti females. J. Insect Physiol. 49, 591–601.

Scott, T.W., Clark, G.G., Lorenz, L.H., Amerasinghe, P.H., Reiter, P., Edman, J.D., 1993. Detection of multiple blood feeding in Aedes aegypti (Diptera: Culicidae) during a single gonotrophic cycle using a histologic technique. J. Med. Entomol. 30, 94–99.

Shaw, P.J., Cirelli, C., Greenspan, R.J., Tononi, G., 2000. Correlates of sleep and waking in Drosophila melanogaster. Science 287, 1834–1837.

Takken, W., van Loon JJ, Adam, W., 2001. Inhibition of host-seeking response and olfactory responsiveness in Anopheles gambiae following blood feeding. J. Insect Physiol. 47, 303–310.

Taylor, B., Jones, M.D., 1969. The circadian rhythm of flight activity in the mosquito Aedes aegypti (L.). The phase-setting effects of light-on and light-off. J. Exp. Biol. 51, 59–70.

Upshur, I.F., Fehlman, M., Parikh, V., Vinauger, C., Lahondère, C., 2023. Sugar feeding by invasive mosquito species on ornamental and wild plants. Sci. Rep. 13, 22121.

Villanueva, J.E., Livelo, C., Trujillo, A.S., Chandran, S., Woodworth, B., Andrade, L., Le, H.D., Manor, U., Panda, S., Melkani, G.C., 2019. Time-restricted feeding restores muscle function in Drosophila models of obesity and circadian-rhythm disruption. Nat. Commun. 10, 2700.

Whiten, S.R., Ray, W.K., Helm, R.F., Adelman, Z.N., 2018. Characterization of the adult Aedes aegypti early midgut peritrophic matrix proteome using LC-MS. PLoS One 13, e0194734.

Wilks, A., Heinzl, G., 2014. Heme oxygenation and the widening paradigm of heme degradation. Arch. Biochem. Biophys. 544, 87–95.

Wynne, N.E., Applebach, E., Chandrasegaran, K., Ajayi, O.M., Chakraborty, S., Bonizzoni, M., Lahondère, C., Benoit, J.B., Vinauger, C., 2024. Aedes albopictus colonies from different geographic origins differ in their sleep and activity levels but not in the time of peak activity. Med. Vet. Entomol. 10.1111/mve.12765

Zheng, X., Yang, Z., Yue, Z., Alvarez, J.D., Sehgal, A., 2007. FOXO and insulin signaling regulate sensitivity of the circadian clock to oxidative stress. Proc. Natl. Acad. Sci. U. S. A. 104, 15899–15904.

Zhou, G., Kohlhepp, P., Geiser, D., Frasquillo, M.D.C., Vazquez-Moreno, L., Winzerling, J.J., 2007. Fate of blood meal iron in mosquitoes. J. Insect Physiol. 53, 1169–1178.

